# The bacterial community structure dynamics in *Meloidogyne incognita* infected roots and its role in worm-microbiome interactions

**DOI:** 10.1101/2020.03.25.007294

**Authors:** Timur Yergaliyev, Rivka Alexander-Shani, Hanna Dimeretz, Shimon Pivonia, David McK. Bird, Shimon Rachmilevitch, Amir Szitenberg

## Abstract

**Background:** Plant parasitic nematodes such as *Meloidogyne incognita* have a complex life cycle, occurring sequentially in various niches of the root and rhizosphere. They are known to form a range of interactions with bacteria and other microorganisms, that can affect their densities and virulence. High throughput sequencing can reveal these interactions in high temporal, and geographic resolutions, although thus far we have only scratched the surface. We have carried out a longitudinal sampling scheme, repeatedly collecting rhizosphere soil, roots, galls and second stage juveniles from 20 plants to provide a high resolution view of bacterial succession in these niches, using 16S rRNA metabarcoding.

**Results:** We find that a structured community develops in the root, in which gall communities diverge from root segments lacking a gall, and that this structure is maintained throughout the crop season. We detail the successional process leading toward this structure, which is driven by interactions with the nematode and later by an increase in bacteria often found in hypoxic and anaerobic environments. We show evidence that this structure may play a role in the nematode’s chemotaxis towards uninfected root segments. Finally, we describe the J2 epibiotic microenvironment as ecologically deterministic, in part, due to active bacterial attraction of second stage juveniles.

**Conclusions:** High density sampling, both temporally and across adjacent microniches, coupled with the power and relative low cost of metabarcoding, has provided us with a high resolution description of our study system. Such an approach can advance our understanding of holobiont ecology. *Meloidogyne* spp., with their relatively low genetic diversity, large geographic range and the simplified agricultural ecosystems they occupy, can serve as a model organism. Additionally, the perspective this approach provides could promote the efforts toward biological control efficacy.

## Introduction

Root knot nematodes (RKN; genus *Meloidogyne*) are among the world’s most devastating plant pathogens, causing substantial yield losses in nearly all major agricultural crops [1]. *M. incognita* and closely related species are found in all regions that have mild winter temperatures [2], and are regarded as one of the most serious threats to agriculture as climate change progresses [3]. In their life-cycle, *M. incognita* hatch in the soil, then invade a root. Thus, the nematodes are exposed to the soil microbiome, rhizobacteria, root epiphytes and endophytes. Once inside the roots, the females modify the cells in order to establish a feeding site and form the characteristic knots for which they are named. Each knot contains at least one nematode feeding from a unique cell-type (the giant cells), surrounded by a gall of dividing cortical cells [4–7]. Throughout its life cycle stages, *M. incognita* are known to interact with microbes, such as cellulase-secreting bacteria and plant effector secreting bacteria, or bacterial and fungal antagonists [8–13]. Consequently, it appears that the geographic or temporal variation in the rhizosphere and root bacterial and fungal communities, can partly explain why such a near isogenic group of nematodes [14, 15] would display variable infestation success [8–11, 16–19].

The interaction of *M. incognita* virulence and microbial taxa in the various niches they occupy, have been studied in the context of biological control, revealing complex relationships, which efficacy diminishes with the transfer from lab to field [20]. Common themes in this line of research include the isolation of *Meloidogyne* pathogens from the cuticle of second stage juveniles (J2) [21–23], or the identification of soil microbes and bacterial volatile compounds with antagonistic effects [24, 25], key taxa including *Rhizobia* [26], *Trichoderma* and *Pseudomonas* [27, 28], *Pasteuria* [29], *Pochonia* [30, 31] and some mycorrhiza [31–33].

Despite the importance of this plant parasite, and its close ties with its cohabiting microbiome, microbial ecology studies utilizing deep sequencing approaches are a handful. Only a few studies have attempted to characterize the taxonomic and functional core microbiota [34, 35], or tie the microbial community composition in the soil or plant with RKN suppressiveness [36–39]. In such studies, temporal dynamics of the microbiome in each of the various niches the nematode occupy at its different life stages, or along the crop season, is not often considered.

In this study we aimed to describe the rhizosphere, root, gall and J2 bacterial succession across the primary nematode life cycle and throughout the crop season to understand microbial interactions as time progresses. Temporal dynamics of the microbiome may have implications for the nematode’s life cycle, but it is also important for the successful application and development of biocontrol agents. To achieve this, we sampled the niches RKN occupy, at six time points, in 20 eggplant plants in Southern Israel.

## Results

To study the temporal dynamics of the bacterial community in the rhizosphere, the roots, galls and J2s in infected eggplants, we sampled these four niches from 20 plants in six time points throughout the crop season, lasting 5 months. For each sample, we sequenced a 16S rRNA metabarcoding library, based on the V3-V4 hypervariable regions, on the Illumina MiSeq platform yielding 34,073,619 sequence read pairs. A curated dataset of 306 samples, containing 10,416 amplicon sequence variants (ASV), was retained following sequence error correction, chimera detection, and the exclusion of organelle sequences, low abundance variants and low abundance samples (see Methods section). This dataset included 150 infected rhizosphere samples, 74 root samples, 61 gall samples, and 21 J2 samples. To describe the succession of bacteria throughout the crop season we always selected the largest galls in the root sample and an adjacent “Infected root” sample lacking a gall. Among the retained samples, read counts ranged between 8,828 and 107,687, with an average read count of 30,145. According to alpha-rarefaction curves (<u>Fig. S1</u>) a sequencing depth of 8,828 was sufficient to capture rare taxa. The bioinformatics analysis carried out for this paper is available as Jupyter notebooks, along with input and output files, in a GitHub repository (https://github.com/DSASC/yergaliyev2020) and on Zenodo (DOI: 10.5281/zenodo.3724182).

### Taxonomic bacterial community composition of the soil, root, gall, and *J2* samples

Bacteria in the system belonged to 37 phyla, with a large variation among niches (Fig. 1). The most abundant phylum was Proteobacteria, followed by Planctomycetess, Actinobacteria, and Chloroflexi. For rhizosphere soil samples, 36 phyla were detected, the most abundant of which being Proteobacteria (27.3%), Planctomycetes (18.3%), Chloroflexi (11.7%), Actinobacteria (11.4%), Firmicutes (7.7%) and Bacteroidetes (6.2%). Of all niches, Proteobacteria had the lowest relative abundance in the soil. Microbiomes of infected root fragments lacking a gall (hereafter “infected roots”), and of gall samples, contained 29 and 30 phyla respectively, and were very similar to each other on the phylum level. In both infected root and gall samples, the most abundant phyla were Proteobacteria (42.8% and 44.5%), Planctomycetes (15.9% and 13.4%), Actinobacteria (12.5% and 10.2%) and Bacteroidetes (7.3% and 10.6%). However, consistent differences between the infected-root and gall samples were observed in Chloroflexi (5% and 4%), Firmicutes (3.4% and 1.9%) and Verrucomicrobia (2.9%and 6.5%).Proteobacteria was the most abundant phylum in J2s as well (74.8%), almost dominating the community. However, unlike the soil, infected-root and gall samples, Bacteroidetes (12.7%) was relatively more abundant than Actinobacteria (3.9%) and Planctomycetes (1.8%). Verrucomicrobia (1.7%), Firmicutes (1.6%), Chloroflexi (1.4%), Patescibacteria (0.7%), Cyanobacteria (0.5%), and Acidobacteria (0.4%) were also observed in J2s.

**Fig. 1:**
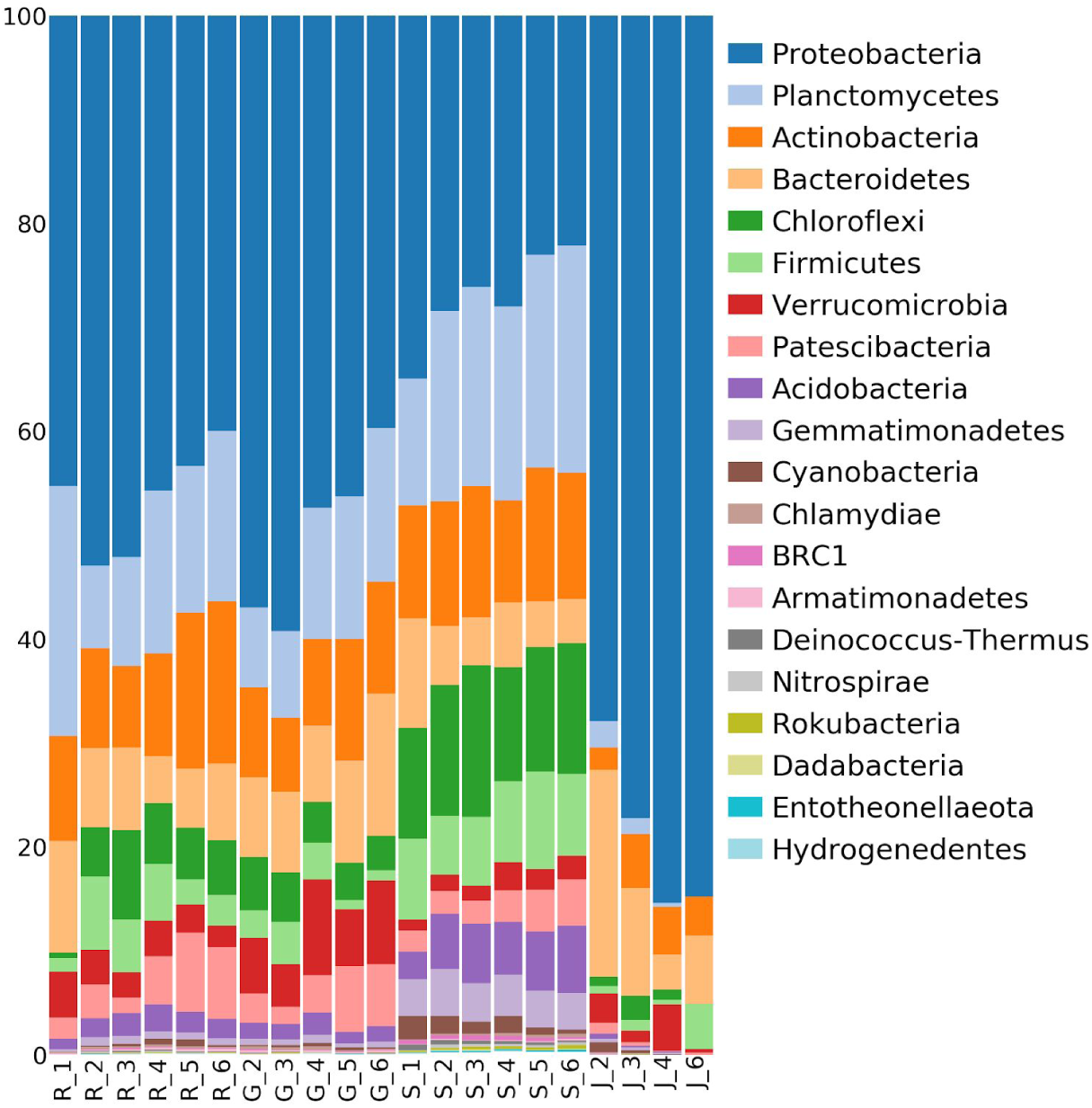
Bacteria relative abundances. Phylum level community compositions (20 most abundant phyla), pooled by niche and time point. R - root section lacking a gall, G - gall, S - rhizosphere soil, J - second stage juvenile (J2). The integers represent time points 1 to 6.

While Proteobacteria were most abundant in J2s and in any of the niches occupied by the nematode, Proteobacteria in the soil, infected-roots and galls were represented mostly by Alphaproteobacteria, while in J2s Gammaproteobacteria (55.4%) dominated the community. Additionally, the relative abundances of Proteobacteria decreased in all niches as time progressed, but not in the J2s, where it increased with time. In contrast, Planctomycetes were the most abundant in the soil (17-22%) and their relative abundance increased with time. Naive root samples taken just prior to planting, demonstrated notably higher relative abundances of Planctomycetes, decreasing after planting. In J2s, unlike other sample types, Planctomycetes relative abundances decreased with time.

### Alpha diversity

To study the temporal changes in alpha-diversity during the vegetation season we calculated the total observed ASV, Pielou’s evenness [40], Shannon’s diversity [41] and Faith’s phylogenetic diversity (Faith’s PD) [42] indices in each sample. We then summarized them by niche at each time-point (Fig. 2A). ASV count increased through the crop season in a moderate fashion in the infected root samples (green dashes), and more steeply in the rhizosphere (solid brown line). This increase did not affect the phylogenetic diversity of ASVs, as it was accompanied by a similar increase in Faith’s PD,m while Shannon’s and Pielou’s indices were only very slightly perturbed. The temporal increase of alpha diversity in the roots as time progressed corresponded with the alpha diversity increase in the rhizosphere.

**Fig. 2:**
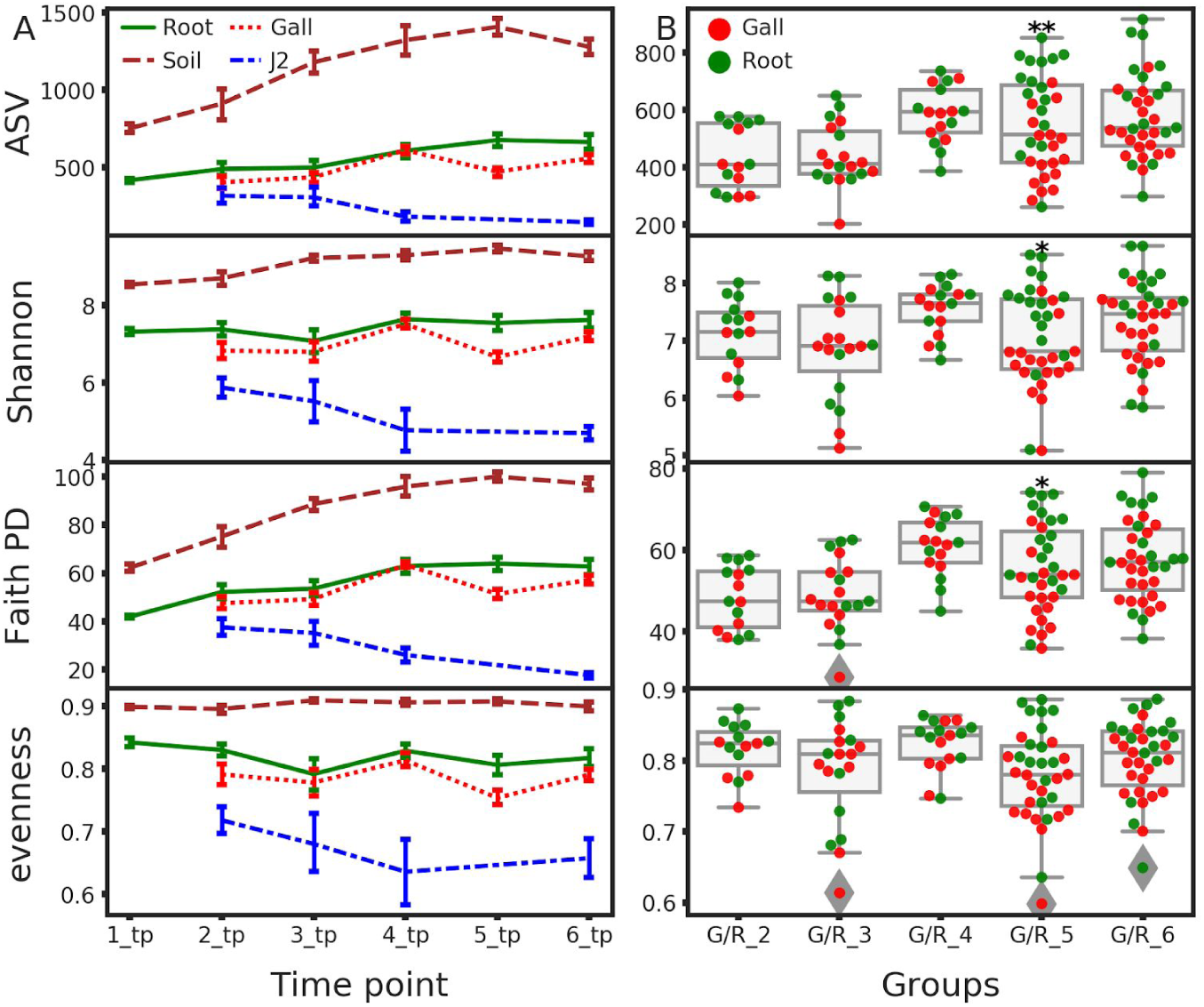
Alpha diversity indices. **A:** A longitudinal representation of alpha diversity indices (Shannon’s diversity index, Faith’s PD, observed ASVs, and evenness), for each niche. The error bars represent the standard deviation. **B:** Alpha diversity indices distribution in gall (G) and infected root (R) samples from time points 2 to 6. Significant differences between galls and roots, according to a Kruskal-Wallis test are indicated by * (q-value < 0.05) or ** (q-value < 0.01).

Alpha diversity in gall samples diverged from that of infected roots toward the end of the crop season. Considering all the metrics, which have decreased in time point 5 in comparison with the infected root samples (Fig. 2B, kruskal-wallis q-value = 0.008, 0.021, and 0.011 for the observed ASVs, Shannon’s diversity index, and Faith’s PD, respectively), this was likely due to an increase in relative abundance of a previously existing and phylogenetically narrow cohort of ASVs in the galls. J2s, collected from root surfaces starting with time point 2, had lower alpha diversity measures than other sample types, and they decreased further through the crop season. The reduction was reflected by all indices, indicating that a phylogenetically narrow group of ASVs gradually took over the J2 cuticule community as time progressed.

### Beta diversity

Beta diversity analyses were performed in order to study temporal and niche effects on the bacterial community composition. Weighted and unweighted pairwise UniFrac distances [43] were computed to account for changes in relative abundances or in the presence and absence of ASVs, respectively. Principal coordinate analysis (PCoA) [44, 45] and biplots [45] were then used to visualize the relationships among the different data classes, and the key ASVs that explain them. ASVs were referred to by both their taxonomic assignment and the first six digits of their MD5 digests, referring to the full digests, as they appear in the representative-sequences fasta file and biom table.

In the PCoA analysis, unweighted (Fig. 3A) and weighted (Fig. 3B) UniFrac distances among the samples were explained by the niche of origin (axis 1, explaining 18.2% and 28.3% of the total variance, for unweighted and weighted UniFrac distances) and by time (axis 2, explaining 8.5% and 13.6% of the total variance). J2 samples were least affected by time and were most similar to early season roots throughout the season, mostly in terms of ASV presence and absence (Fig. 3A). Nevertheless, they were distinctly different from the root sample communities, suggesting that the composition of the nematode-associated bacterial community is quite unique and notably different from other niches. Biplot results revealed an increase in the relative abundance of ASVs assigned to *Pseudomonas* in J2s from all time points, and early season roots, compared to other sample classes (Fig. 3B) and the increase of ASVs assigned to *Azospirillum*, and *Reinheimera*.

**Fig. 3:**
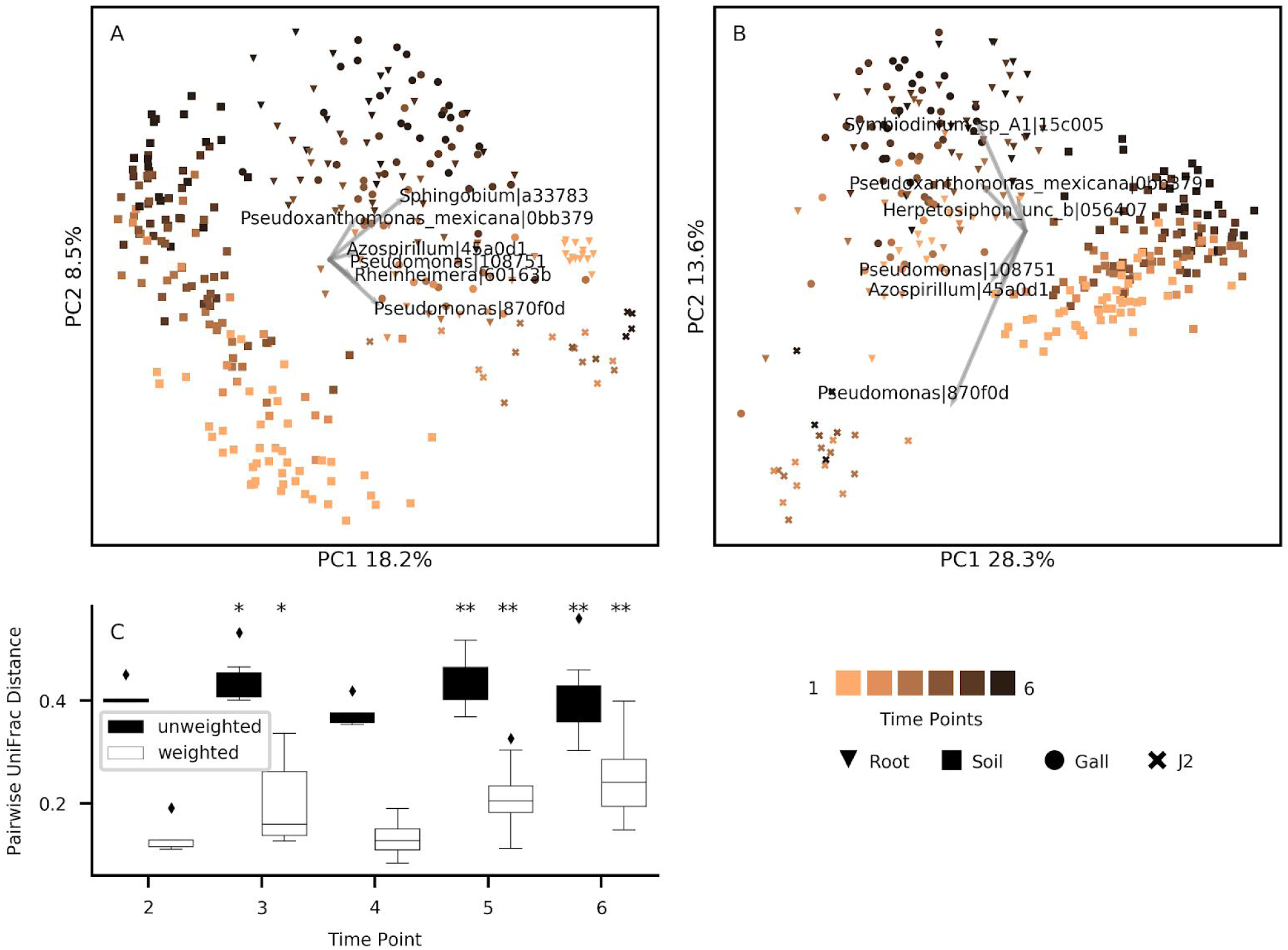
UniFrac distances among samples. **A:** Unweighted and **B:** weighted UniFrac PCoA ordinations and biplots, and **C:** the distribution of pairwise UniFrac distances between infected root and gall samples in time point 2 to 6. The markers represent the different niches and time points. Explanatory ASVs are denoted by the genus they were assigned to and the first six characters of their MD5 digest. Asterisks denote significant Wilcoxon test results. *: q-value < 0.05. **: q-value < 0.01.

Gall samples (sphere markers; Fig. 3) differentiated from root (triangle markers; Fig. 3) communities in late season time points, but with some overlap between the two niches. To test whether the observed differentiation of infected root and gall samples as the season progressed was significant, we carried out pairwise Wilcoxon paired tests [46] and Permanova tests [47]. For time point 5 and 6, the distance between the sample types was significantly larger than zero (q-value < 0.01), for both tests and both distance measures, but more discernible for the weighted UniFrac distances (Fig. 3C). The weighted UniFrac distance between root and galls samples was significantly larger than zero at time point 4 as well (q=0.024), when tested with Permanova. The stronger signal received from the weighted UniFrac distance indicates that this divergence is mainly due to few bacteria, which took over the gall communities, and less so due to the introduction of new bacteria.

To identify the ASVs responsible for the differentiation between gall and infected root samples, we repeated the PCoA and Biplot analyses, including only galls and infected root samples from the last time point (Fig. 4). Using the weighted UniFrac distance matrix, the combination of axes 1 and 2 segregated the two niches almost completely with 19.7% and 15.0% of the variance explained. Key ASVs responsible for this segregation belonged to Planctomycetes, *Pseudoxanthomonas* spp., genera from the Rhizobium clade (*Allorhizobium, Neorhizobium, Pararhizobium* and *Rhizobium, sensu* Mousavi *et al*. [48], hereafter A/N/P-Rhizobium), *Lechevalieria, Luteolibacter, Pseudomonas* and Saccharimonadales. Only *Pseudomonas* had higher relative abundance in the root samples, while the rest of the ASVs had higher relative abundance in the gall samples.

**Fig. 4:**
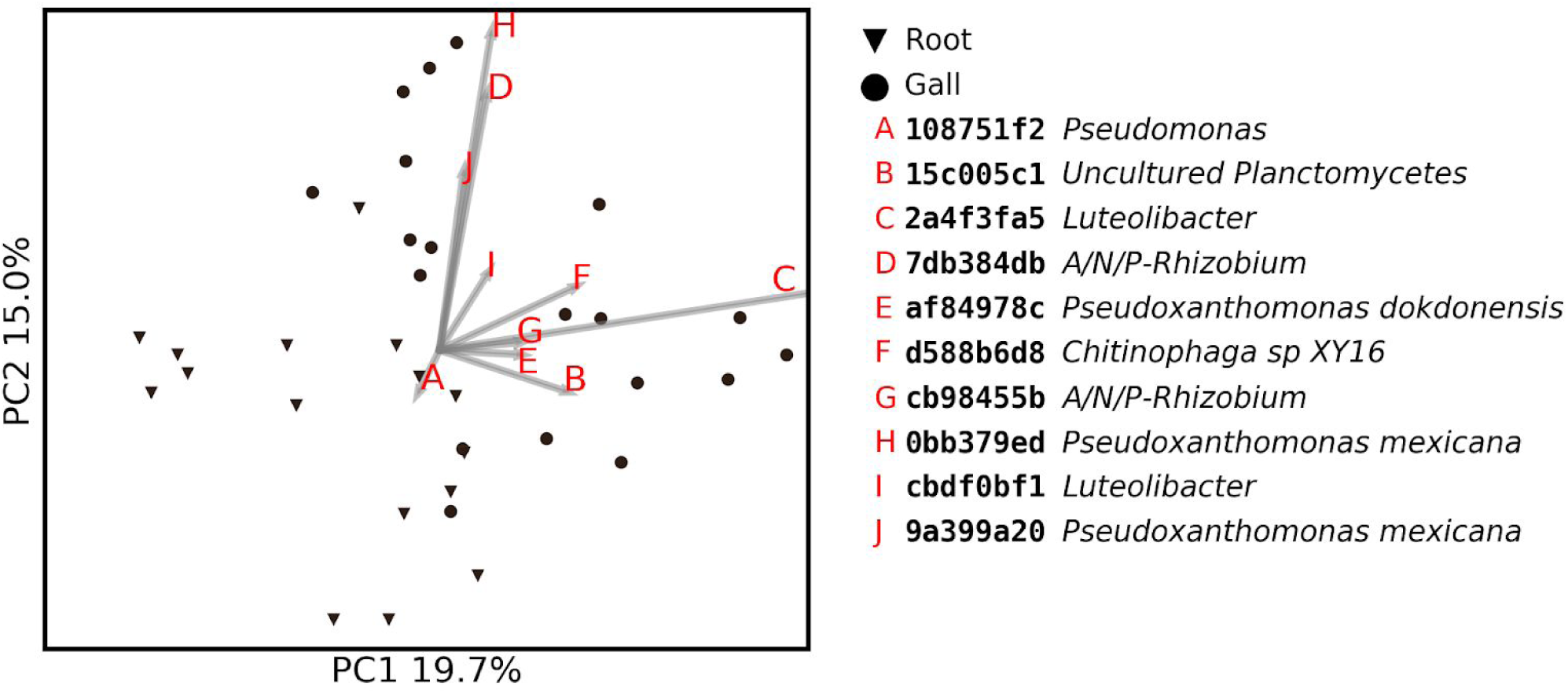
Weighted UniFrac PCoA and biplot for infected root and gall samples from time point 6. Marker shapes represent the niches. The 10 most important explanatory ASVs are represented by the first eight characters of their MD5 digest and their lowest identified taxonomic level.

### Co-occurrence of congeneric core ASVs and their phylogenetic relationships

We treated the ratio between the core ASVs and the core taxa of each niche as a relative ecological-drift measure. A core genus without a respective core ASV is represented by different ASVs in different samples of the niche, indicating that the ecological constraints favoring one species or strain over another are not strong. The larger the deficit in core ASVs compared to the core taxa they are assigned to, the lower the ratio and the larger the drift. We based this approach on the notion that ecological drift can increase the genetic diversity among samples that were obtained from one niche [49]. We formulated the difference as the ratio of core ASV count to core taxa count (ASV to taxa ratio - *R*).

With this notion in mind, among all the niches, only J2 had *R* > 1 on average, in their 100% core microbiome (Fig. 5A). The *R* value in J2, was significantly different from *R* in other niches (q-value < 0.023), in which the ASV count was lower than the taxa count. Additionally, *R* in the infected root samples was also significantly larger than *R* in the soil (q-value = 0.023). When considering only the unique taxa in each niche (Fig. 5B), galls and infected roots had *R* > 1 as well, significantly larger than the soil *R* value (q-value < 0.023). Consequently, J2s appear to provide a more selective microenvironment than the other niches, but the root and gall microenvironments have an intermediate level of stochasticity, between the soil and J2 niches.

**Fig. 5:**
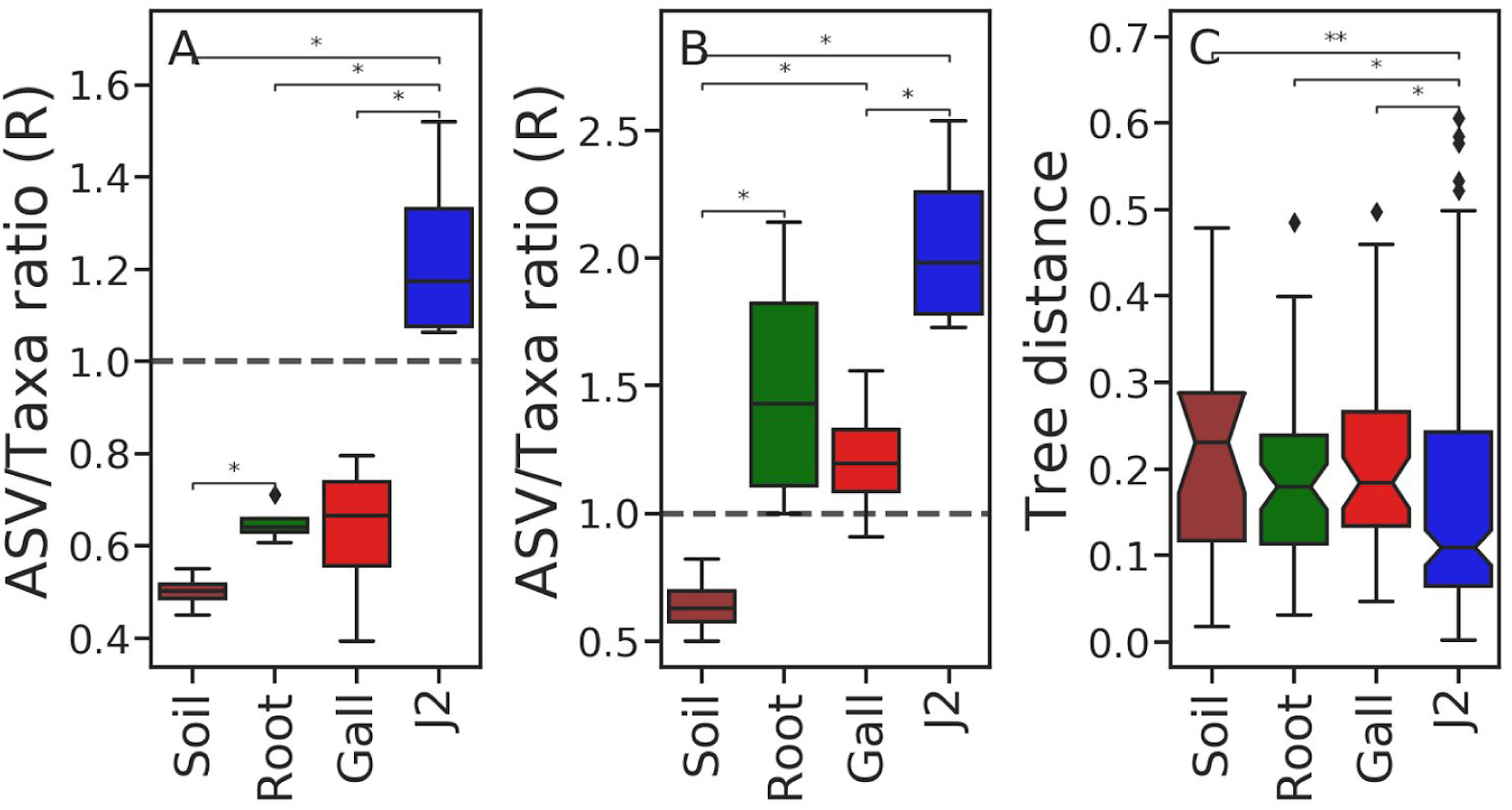
Comparative ecological drift in the sampled niches. The ratio - *R* between (**A**) all core ASV count and all core taxa count, or between (**B**) the unique core ASV count and unique core taxa count, as a relative measure of ecological drift in each niche. The distribution of pairwise patristic distances between congeneric ASVs (**C**) was also used to compare stochasticity among the niches. Only time point 2, 3, 4 and 6 were used in all analyses.

We have taken an additional, phylogenetic approach, towards comparing the strength of any deterministic forces shaping the communities in the various niches. For each niche, we computed the pairwise patristic distances between core ASVs sharing a taxon assignment. We expected a very selective niche to sustain congeneric ASVs that are more closely related to one another than congeneric ASVs in a more stochastic niche (Fig. 5C). J2 had a significantly lower median partistic distance than other niches (q-value < 0.014) indicating that congeneric ASVs in the J2 samples are more closely related than congeneric ASVs in other niches.

### Bacterial succession

In addition to the temporal dynamics of alpha and beta diversity, we investigated the temporal change of discrete features (ASVs or taxa) to characterise the bacterial succession in various niches. We focused our investigation on features that we have identified as “important” or “dynamic” (see Methods section), based on a features volatility analysis [50]. We also investigated the two most abundant ASVs belonging to the included taxa, were they not already considered. The mean relative abundance of features is presented as a heatmap (Fig. 6), organised according to niche (Fig. 6A-C) and time point. Blue shades represent the relative abundance of each taxon across the time points, green shades represent the relative abundance of each ASV within the pool of ASVs assigned to a certain taxon, and the purple shades represent the temporal distribution of the ASV or taxon. Dashed borders separate a taxon and the ASVs assigned to it.

**Fig. 6:**
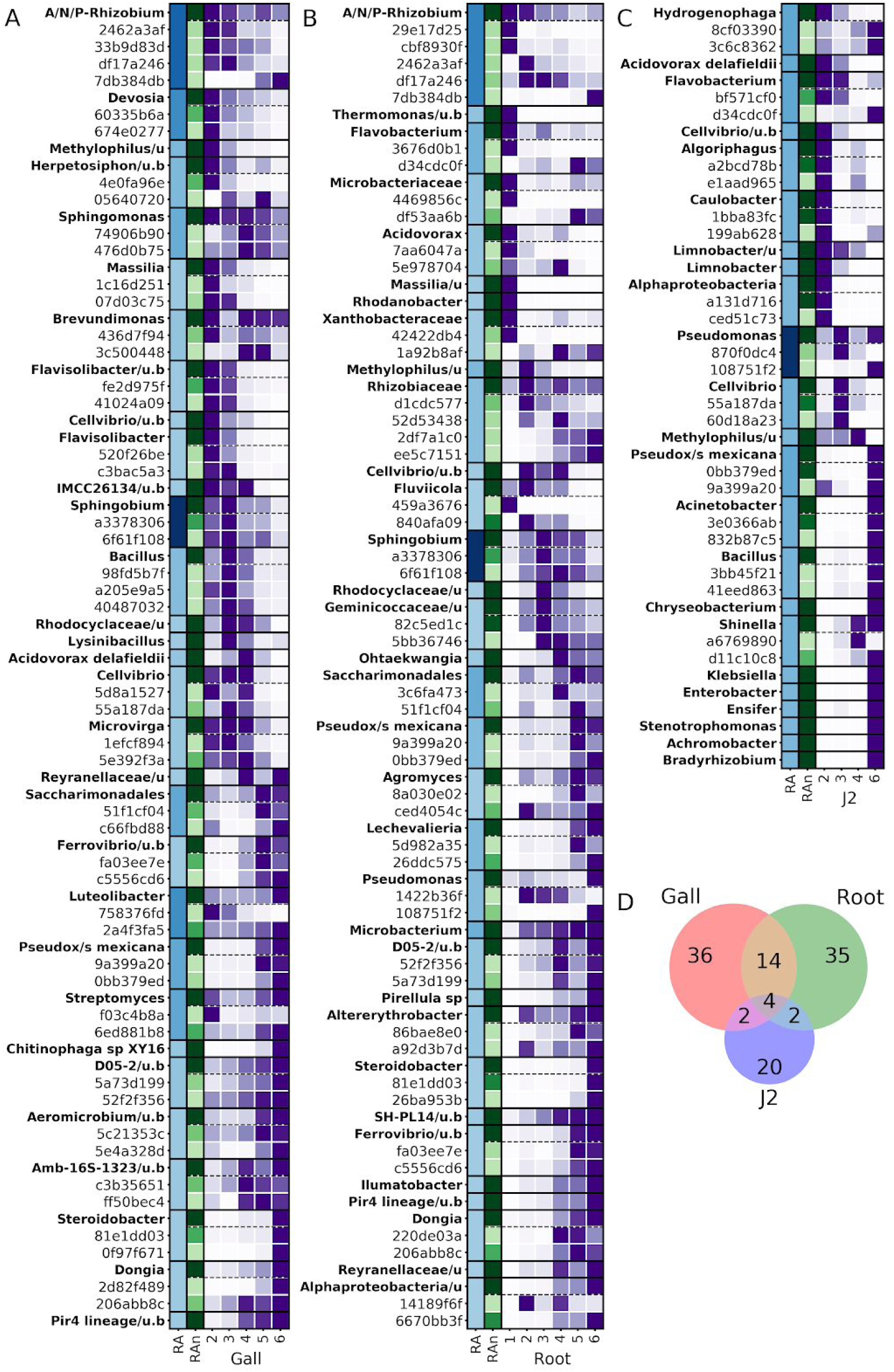
Bacterial succession of dynamic and important features. The most important and dynamic features were detected using a feature volatility analysis (see Methods section). The curated cohort was selected separately for the galls (**A**), infected roots (**B**) and J2 (**C**), and include taxa (bold font) and some of their ASVs. The relative abundance of each taxon is indicated in blue (*RA*). The relative abundance of each ASV, among the ASVs assigned to a single taxon, is indicated in green (*RAn*), normalised by the relative abundance of the taxon. The temporal distribution of each feature is indicated in purple (time points), normalised by the peak relative abundance of the feature. The number of shared and unique ASVs in the curated list of each niche is indicated in the Venn diagram (D).

A comparison of the three heatmaps reveals largely independent sets of bacteria in each niche, which most explain temporal changes, as illustrated by the Venn diagram of the heatmap ASVs (Fig. 6D). Within the galls (Fig. 6A), an “early” bacterial community, including 14 taxa (A/N/P-Rhizobium, *Devosia, Methylophilus, Herpetosiphon, Sphingomonas, Massilia, Brevundimonas, Flavisolibacter, Cellvibrio*, the uncultured Verrucomicrobia IMCC26134, *Sphingobium, Bacillus*, Rhodocyclaceae, *Acidovorax delafieldii* and *Microvirga*), is gradually displaced by a “late” gall community including 12 taxa (Reyranellaceae, Saccharimonadales, *Ferrovibrio, Luteolibacter, Pseudoxanthomonas mexicana, Streptomyces, Chitinophaga, Aeromicrobium*, the uncultured Rhizobiales Amb-16S-1323, *Steroidobacter, Dongia* and uncultured Planctomycetaceae sp. belonging to lineage Pir4). This displacement already begins and intensifies in time points two and three, within the primary nematode life cycle.

Another important aspect of the gall bacterial succession is the origin of taxa. Only a few taxa in the root clearly originated from the naive roots. In the early community, these include *Methylophilus, Sphingobium and Acidovorax delafieldii* (<u>Fig. S2</u>). Of these, only *Sphingobium* persisted successfully throughout most of the crop season. Conversely, most of the genera detected in the galls emerged from the soil, and were sometimes displaced by congenerics in later time points. Most notably, A/N/P-Rhizobium were already represented in the naive roots, but then were effectively competed against by soil congenerics, which then shared the gall A/N/P-Rhizobium community (e.g. root originated ASV cbf5930fe vs. soil originated ASV 33b9d83d1; <u>Fig. S2</u>). In the late gall community, only *Luteolibacter* clearly emerged from the naive roots and not from the soil (<u>Fig. S2</u>). *Luteolibacter* represented ∼7% of the gall community by the end of the season. ASV 2a4f3fa50 outcompeted other congeneric ASVs to monopolize the *Luteolibacter* community.

### Attraction assay

In 2017, gall bearing eggplant roots were sliced and washed with PBS in order to attempt the isolation of RKN related bacteria (supplementary file S1). As RKN pathogens often actively attract J2s, we carried out an attraction assay, testing the attraction of J2s to each of two isolates given the isolate and fresh root as options, or the sterile medium and a fresh root as control. One isolate in particular presented a 10-fold larger attraction of J2s than the root (supplementary file S1). Upon sanger sequencing (supplementary file S1), this isolate had an identical V3-V4 region sequence as *Pseudomonas* 108751f2 (Fig. 4).

To test whether compounds secreted by this isolate can attract J2s, we carried out an additional attraction assay, in which J2s were allowed to select between a fresh root fragment and one of the following: whole bacterial medium filtrate, containing the isolate’s exudates, the < 3 kDa fraction of this filtrate, the 3-100 kDa fraction of the filtrate and the fraction of compounds larger than 100 kDa (see Methods section). The results are summarised in Fig. 7, showing a significantly larger attraction of J2s to the < 3kDa fraction of the filtrate, than to the other fractions or the whole filtrate (0.0009 < q-value < 0.027). Upon observation under a dissecting microscope, J2 nematodes in the < 3 kDa fraction were active for at least three days. In the 3-100 kDa fraction, nematodes were only mildly active after three days. The few individuals found in the >100 kDa fraction were inactive after 2 hours.

**Fig. 7:**
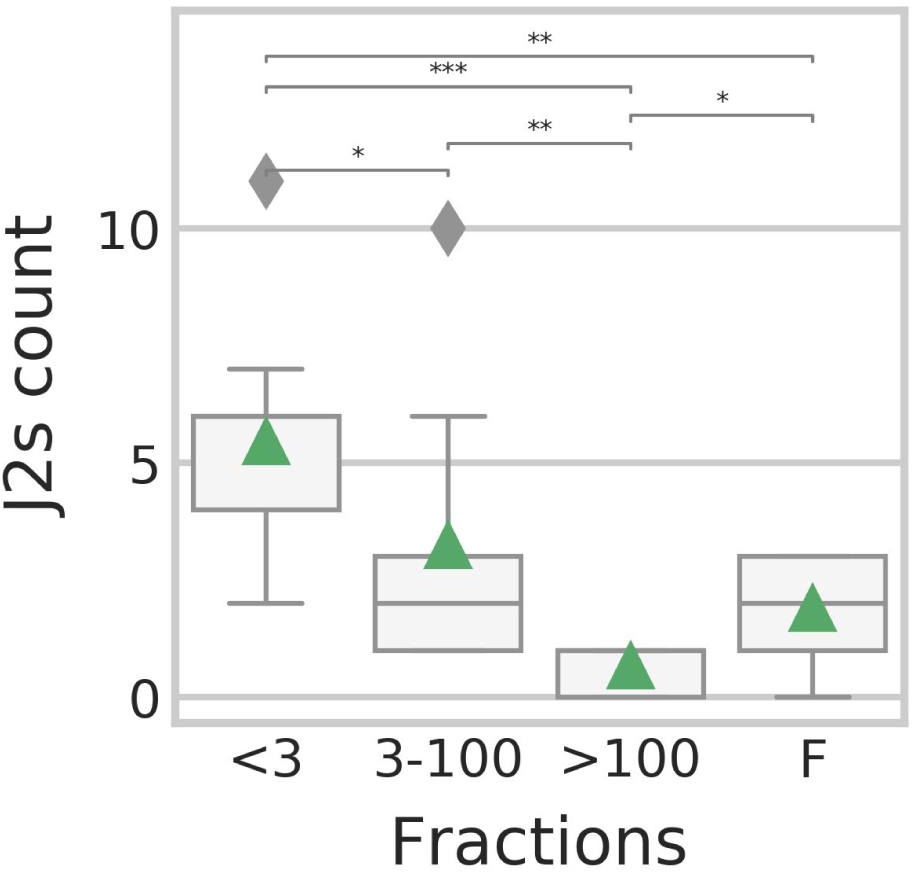
Attraction assay. The number of J2 nematodes attraction to a *Pseudomonas* filtrate, when presented with both the filtrate and a root fragment. X axis: the size of compounds in the filtrate (kDa). F - whole filtrate. Y-axis, J2 nematode count. Asterisks indicate significant differences according to Mann-Whitney U test. ***: q-value < 0.001. **: q-value < 0.01. *: q-value < 0.05.

## Discussion

### The root community structure at the end of the crop season

Animals and plants host a large number of microbes, carrying out many interactions, which can modulate the functions and behaviours within the complex. This microbial community, like any other ecological community, is subject to succession. In this study, we investigated the structure of the bacterial community in RKN infected roots and its interaction with the communities in the soil and J2 nematodes, with respect to the community dynamics along the nematode’s primary life cycle and the crop season. According to beta-diversity indices (Fig. 4), gall communities diverge from those of adjacent root sections lacking a gall, late in the crop season, revealing a highly structured root community. According to alpha (Fig. 2B) and beta diversity (Fig. 3C) pairwise test, this structure starts to develop even earlier in the season.

The mature gall community differs from that of adjacent root segments by the higher relative abundance of bacteria with known nematostatic (*Pseudoxanthomonas* spp.; [51]) and antibiotic activity (*Lechevalieria* sp. [52–55]), RKN symbionts participating in the structural modification of the root (A/N/P-Rhizobium [13]), nematode egg-shell feeding bacteria (*Chitinophaga*; [56, 57]), or bacteria providing protection from reactive oxygen species (ROS) (*Luteolibacter*; [58]). Conversely, the adjacent root segments present a higher relative abundance of a *Pseudomonas* sp. (ASV 108751f2), which increase occurred late in the season (Fig. 6B). This *Pseudomonas* sp. was isolated and experimentally proven to be highly attractive to the J2, as well as the fraction of its exudates, which contains molecules smaller than 3 kDa. It is thus possible that this structured bacterial community we have identified plays a role in providing J2s in the soil with the chemical cues they require to identify the healthier sections of the already deteriorated root system. Additionally, some endophytic *Pseudomonas* spp. have been shown to degrade ROS, and can thus further facilitate the repeated infection of parasitic nematodes [59, 60]. The same *Pseudomonas* sp. strain (ASV 108751f2) was also detected on J2s in increasing densities at the end of the crop season (Fig. 6C). This could be an artifact of the Baermann tray method, but the evidence supporting J2’s active attraction to this isolate could point to a more elaborate form of interaction in which the bacterium first attracts the nematode to hitchhike into the root, and once established, can help guide the nematode into less abstracted sections of the root. Possibly, the benefit to the J2 is even more immediate, if this *Pseudomonas* sp. provides a low-ROS microenvironment to the J2, upon its penetration to the root. Such a relationship has already been recorded between *M. hapla* and an epibiotic *Microbacterium [61]*.

A/N/P-Rhizobium, another taxon which occurred both in the gall and J2 samples, showed a covarying pattern of relative abundance between the two niches (<u>Fig. S2</u>). It has been shown in the past that bacteria belonging to this group were able to successfully invade plant hosts by using nematodes as a transicion vector [62]. Therefore, this observed covarying pattern of relative abundance may be non-incidental.

### The bacterial succession in the gall along the primary RKN life cycle and the crop season

Upon consideration of the most dynamic fractions of the bacterial communities, the processes leading to the end result described above are exposed as a gradual community shift (Fig. 6A). The early gall community comprises bacteria which confer structural modifications to the root (A/N/P-Rhizobium [13], *Devosia* sp. [63], and then *Microvirga* [64]), fixing nitrogen (A/N/P-Rhizobium [65], *Devosia* sp. [63, 66, 67], *Microvirga* [64], *Sphingomonas* [68] and later *Cellvibrio [69], Microvirga* [64] and possibly *Bacillus [70]*) and bacteria capable of degrading polysaccharides (*Herpetosiphon* [71, 72], *Massilia* [73], *Verrucomicrobia [74, 75], Cellvibrio [76], Sphingobium* [77] and possibly *Bacillus [78]*). While some potential chitin feeders form a part of this community early on (*Herpetosiphon* [79, 80], *Sphingomonas* [81] and *Massilia [73]*), time point 3, 40 days from planting and onwards, sees a gradual addition of chitin feeders (*Cellvibrio [82], Lysinibacillus* [83], *Streptomyces* [84] and possibly *Bacillus [83, 85]*), possibly due to an increase in egg-shell density.

Since we sampled the largest galls at each time point, the late season galls (time points 4-6) likely represent already inactive galls in most cases. The most striking characteristic of the late gall community, is a rapid increase in bacteria capable of anaerobic or hypoxic growth. Only ASVs of *Sphingomonas*, known to include facultative anaerobic species [86], persisted throughout the season. Even a Rhizobiales representative which occurs in the late gall community (Amb-16S-1323) seems to occur in hypoxic and anaerobic environments, unlike the early season representatives of the order which are aerobic, plant related bacteria (<u>Fig. S3</u>). Eight other genera, which ASVs increased in this time frame have known anaerobic or hypoxic species (*Bacillus* [87, 88], *Rhodocyclaceae [89], Acidovorax delafieldii [90], Ferrovibrio [91], Pseudoxanthomonas mexicana[92], Chitinophaga [56], Steroidobacter* [93] and a Pir4 lineage bacterium [94]). Lastly, some of the taxa comprising the late gall community have been associated with plant parasitic nematode soil suppressiveness or RKN antagonism (*Chitinophaga* and *Streptomyces [95–99]*)

### The second stage juveniles epibiotic microbiome

Almost all bacteria that were found to be most important or dynamic in second stage juveniles (Fig. 6C) are associated with RKN control in the literature [100–106], although *Cellvibrio* may be assisting their root penetration [107]. While the bacterial communities of the soil and root samples consistently shifted in time, according to UniFrac pairwise distances (Fig. 3A & B), the J2 community beta-diversity indices remained relatively very constant. Furthermore, most of the core genera in the J2 community were represented by the same ASVs in all samples and time points, unlike other niches, and co-occurring congeneric ASVs were more closely related in J2 samples than in other niches (Fig. 5). Presumably, the J2 cuticule is a very selective environment and specialist bacteria can predictably outcompete other bacteria that exist in the root or soil. It is possible that like in many other organisms, in which skin epibionts play a role in preventing pathogens from settling [108], these J2 epibionts may have to evade antimicrobial mechanisms exerted by the host. Concomitantly, some bacteria [16, e.g., 109] actively attract or attach to J2 nematodes and colonise their surface in various forms of symbiosis. The *Pseudomonas* sp. isolate which density increased in the J2 samples at the end of the crop season (Fig. 6C) and was shown to attract the J2 nematodes (Fig. 7) would also belong to this group of J2 symbionts. This is another mechanism maintaining the low genetic diversity of the J2 epibionts, which even reduces alpha-diversity with time (Fig. 2A). Although the Baermann tray extraction method would potentially bias the result by contaminating the perceived J2 bacterial community with root and soil bacteria, we believe this effect was small, given the fact that the J2 microbiome remained constant, in spite of the very dynamic community in the other niches.

## Conclusion

In this study, *via* a longitudinal approach and a large number of replicates, we were able to robustly describe a bacterial community structure within *M. incognita* infected roots. This structure seems to be preserved even at the very end of the crop season and to potentially play a role in the life cycle of the nematodes. While in the scope of the crop season, there is an increase in bacteria often found in hypoxic and anaerobic environments, some connection to the nematode’s life cycle was also observed, in particularly with the creation of root structural modifications, polysaccharide metabolism and chitin metabolism. With their large geographic range [2, 110], the relatively simplified agricultural ecosystems they occupy and their minimal genetic diversity [14, 15], *M. incognita* could be used as a much needed model organism for terrestrial holobiont ecology studies. Understanding the contribution of variations in the soil and host-crop microbial seed bank on the root community structure and its interactions with environmental factors, could serve our understanding of ecological processes in terrestrial holobionts and the efficacy of biocontrol agents in field conditions [111].

## Methods

### Sampling

Samples were collected from greenhouse cultivated eggplant plants, infected by *M. incognita*, in Hatzeva (Israel). Twenty plants were repeatedly visited through the crop season. Two niches, a root fragment and the rhizosphere soil, were sampled from each plant at each time-point. Additionally, galls and second stage juveniles (J2) were collected when available. In time point 1, root sections were collected prior to planting, to represent the preexisting root endophytic community. To represent the succession of bacteria in the galls through the RKN life cycle and crop season, the largest galls were collected at each time point, and an adjacent 2 cm root fragment, referred to as “infected roots” throughout the manuscript. Roots were also collected from each plant in order to extract J2, using the Baermann tray method, following Williamson and Čepulytė [112]. The J2 were then filtered onto 47mm diameter and 0.45µm pore size cellulose ester filters (GE Healthcare Whatman). All samples were kept at -80 °C until DNA extraction. Table 1 summarizes the number of samples available for analysis after sequencing and data filtration (see below). Rhizosphere soil samples were collected from additional plants and thus the number of soil samples exceeds 20 in most time points. The number of J2 samples in Table 1 was affected mostly by the J2 density in the samples, but also by subsequent data curation (see below).

**Table 1.** Sample collection.

| Time-point | Date | Amount of samples |  |  |  |
| --- | --- | --- | --- | --- | --- |
|  |  | Soil | Root | Gall | J2 |
| 1 | 20.12.17 | 44 | 14* | - | - |
| 2 | 14.01.18 | 16 | 9 | 6 | 8 |
| 3 | 30.01.18 | 21 | 9 | 10 | 7 |
| 4 | 25.02.18 | 26 | 9 | 8 | 2 |
| 5 | 26.03.18 | 24 | 17 | 18 | - |
| 6 | 22.05.18 | 19 | 16 | 19 | 4 |

\* Uninfected roots collected prior to planting

### DNA extraction and 16S rRNA library preparation

The roots were washed with 1% sodium hypochlorite solution and rinsed with distilled water, then a gall and adjacent root fragment were dissected. Each segment was cut into small pieces and placed in a well within a 96 well sample-plate. For each rhizosphere soil sample, 0.25 g sample-plate wells. Root, gall and soil DNA was extracted using the DNeasy PowerSoil kit (Qiagen) following the manufacturer’s instructions. J2 DNA was extracted from the filters using the PowerWater DNA extraction kit (Qiagen), following the manufacturer’s instructions. Metabarcoding libraries were prepared as we previously described [113], using a two step PCR protocol. For the first PCR reaction, the V3-V4 16S rRNA region [114] was amplified using the forward primer ‘5-tcgtcggcagcgtcagatgtgtataagagacagCCTACG GGNGGCWGCAG-’3 and the reverse primer ‘5-gtctcgtgggctcggagatgtgtataagagacagGACTAC HVGGGTATCTAATCC-’3, along with artificial overhang sequences (lowercase). In the second PCR reaction, sample specific barcode sequences and Illumina flow cell adapters were attached, using the forward primer ‘5-AATGATACGGCGACCACCGAGATCTACACt cgtcggcagcgtcagatgtgtataagagacag-’3 and the reverse primer ‘5-CAAGCAGAAGACGGCATACGAGATXXXXX Xgtctcgtgggctcgg-’3’, including Illumina adapters (uppercase), overhang complementary sequences (lowercase), and sample specific DNA barcodes (‘X’ sequence). The PCR reactions were carried out in triplicate, with the KAPA HiFi HotStart ReadyMix PCR Kit (KAPA biosystems), in a volume of 25 µl, including 2 µl of DNA template and following the manufacturer’s instructions. The first PCR reaction started with a denaturation step of 3 minutes at 95 °C, followed by 30 cycles of 20 seconds denaturation at 98 °C, 15 seconds of annealing at 55 °C and 7 seconds polymerization at 72 °C. The reaction was finalized with another one minute long polymerization step. The second PCR reaction was carried out in a volume of 25 µl as well, with 2 µl of the PCR1 product as DNA template. It started with a denaturation step of 3 minutes at 95 °C, followed by 8 cycles of 20 seconds denaturation at 98 °C, 15 seconds of annealing at 55 °C and 7 seconds polymerization at 72 °C. The second PCR reaction was also finalized with another one minute long polymerization step. The first and second PCR reaction products were purified using AMPure XP PCR product cleanup and size selection kit (Beckman Coulter), following the manufacturer’s instructions, and sequenced on an Illumina MiSeq to produce 250 base-pair paired-end sequence reads. The sequencing was carried out by the Nancy and Stephen Grand Israel National Center for Personalized Medicine, The Weizmann Institute of Science.

## Bioinformatics

### Data processing, taxonomy assignment and biodiversity analysis

All the analysis carried out for this paper is available as Jupyter notebooks in a github repository (github: https://github.com/DSASC/yergaliyev2020; Zenodo: DOI: 10.5281/zenodo.3724182), along with the sequence data, intermediate and output files. The bioinformatics analysis was carried out with Qiime2 (v.2019.4/10) [115]. Forward and reverse PCR primers were removed from the MiSeqs reads by q2-cutadapt plugin [116]. Using the q2-DADA2 plugin [117], paired reads were truncated to the length of 267 and 238 bp, for the forward and reverse reads, respectively, and the fist two bases were removed as well. Then the reads were quality-filtered, error corrected, dereplicated and merged. Finally chimeric sequences were removed, to produce the amplicon sequence variants (ASV). For taxonomic assignment, a naive Bayes classifier was trained using taxonomy assigned reference from Silva SSU-rRNA database (v.132, 16S 99%) [118]. Reference sequences were trimmed to the V3-V4 fragment. All ASVs that were identified as mitochondrial or chloroplast sequences, or that were assigned only to Bacteria level, as well as completely unassigned sequences, were filtered out from the feature table. The ASV biom table was further filtered to exclude samples with less than 8828 sequences. An ASV phylogenetic tree was built with the q2-phylogeny plugin, implementing MAFFT 7.3 [119] for sequence alignment, and FastTree 2.1 [120], with the default masking options. Microbial diversity was estimated based on the number of ASVs observed, Pielou’s evenness [40], Shannon’s diversity indices [41] and Faith’s phylogenetic diversity [42] for alpha diversity and weighted and unweighted UniFrac distance [43] matrices for beta diversity. Ordination of the beta-diversity distance was carried out with a principal coordinates analysis (PCoA), and the key taxa explaining beta diversity was obtained with a biplot analysis [44, 45]. Tests for significant differences in alpha and beta diversity between groups of samples representing one niche and one time point were implemented with Kruskal-Wallis test (alpha diversity) [121] or the Wilcoxon [46] and PERMANOVA [47] test (beta diversity). We focused our attention on the gall and infected root sample types. P-values were corrected for multiple testing using the Benjamini-Hochberg procedure [122]. Corrected p-values are referred to as q-values throughout the text. In the Qiime2 environment, the MD5 message-digest algorithm is used to name ASVs by their sequence digest. To make these digests more human readable, we prefixed each digest with the lowest available taxonomic level and the “|” symbol. This change was consistently implemented in the biom table, the representative-sequences fasta file and the taxonomy assignment table.

### Longitudinal feature volatility analysis

The Qiime2 feature volatility longitudinal analysis [50] was carried out to study the temporal changes in relative abundances of each ASV and taxon, separately in the galls, roots and J2. Both ASVs and taxa were analysed, assuming that in some cases discrete ASVs that represent functionally and taxonomically similar bacteria are ecologically interchangeable. In such cases, patterns that would be observed at the species or genus level, might be lost at the ASV level, and *vise-versa*. We grouped ASVs into taxa based on their lowest available taxonomic level. One parameter with which we identified key ASVs and taxa in the system was “importance”. Importance is defined as the euclidean distance of the relative abundance vector of a given taxon or ASV from a null vector of the same length [50]. For each niche, we identified the 15 most important ASVs and 15 most important taxa. We additionally included the 10 ASVs and 10 taxa with the highest net average change among time points (five increasing and five decreasing). Lastly, we included the two most abundant ASVs of the included taxa, were they not already represented. Additionally, for each ASV the list we added the taxon it was assigned to, were it not already included. Fig. 6 summarizes the results of this analysis as a heatmap. It additionally includes a Venn diagram of the ASVs that are represented in the heatmap.

### Core ASV/Taxa ratios and tree distances

To evaluate the relative strength of ecological drift among niches, we computed ASV to Taxa ratios (*R*), within the core microbiome and unique microbiome of each niche and time point. Because ecological drift is expected to increase the genetic diversity among samples [49], under strong drift, similar taxa would be expected to be represented by different ASVs in different samples. Considering two extremities on a range of theoretical possibilities, where a niche is deterministic, congeneric bacteria would be closely related due to shared specialties. However, with strong ecological drift and with niche constraints relaxed, congeneric ASVs would be allowed to diverge.

Core ASV and core taxa were determined by the Qiime2 feature-table plugin separately for several subsets, each containing samples of one niche and one time point, out of the second, third, fourth and sixth time points. Only ASVs or taxa that were detected in all the samples of a given subset were included (100% core microbiome). Another dataset containing only the core ASVs and taxa that were unique to each niche was produced. *R* values were computed for each subset, for the core microbiome and the unique microbiome. The distribution of core and unique *R* values are presented in Fig. 5A & B, respectively. Pairwise comparisons between niches were tested with Mann Whitney U tests [123], corrected for multiple testing using the Benjamini-Hochberg procedure [122].

Since we suspected that the *R* values might be influenced by differences in sample sizes among the subsets, we repeated the analysis with a normalised sample size. To normalise the sample size, we constrained the number of samples in each time point. To do that, we identified the niche with the lowest number of samples in each time point and reduced the number of samples in the remaining niches to conform with that number. The resulting *R* value distributions and the pairwise Mann whitney U tests were similar to those obtained with the full dataset.

A genetic signature of congeneric ASV divergence would also be recorded in their pairwise phylogenetic distances. For each group of congeneric ASVs in the core microbiome, a phylogenetic tree was reconstructed, from which we obtained pairwise patristic distances of congeneric ASVs. Then, the distribution of congeneric ASV patristic distances was computed for each subset and presented as Fig. 5C. The phylogenetic trees were reconstructed as follows. For each genus level taxon, we produced a fasta file of ASVs. The 50 best matches in the Silva 138 16S rRNA database [118] were identified for each ASV, using blastn 2.9.0+ [124]. The ASV and reference sequences were aligned using the L-ins-i algorithm implemented in MAFFT 7.3 [125] and trimmed by trimAl [126], with a 0.1 gap threshold. Maximum likelihood trees were constructed with RAxML 8.2 [127] using the GTR-Gamma model of sequence evolution. Pairwise comparisons of patristic distance distributions were teste with Mann-Whitney U tests [123], corrected for multiple testing with Benjamini-Hochberg Benjamini-Hochberg procedure [122].

### Isolation sources of early and late gall community Rhizobiales

The following steps were taken to summarise the isolation sources of three Rhizobiales taxa from the gall community, including the group of uncultured Rhizobiales Amb-16S-1323, *Devosia* spp. and the A/N/P-Rhizobium cluster. For each taxon, all the sequences available on Silva [118] were downloaded as fasta files, and their GenBank entries were retrieved based on their accession number, using the BioPython Entrez python module [128]. Isolation sources, taken from the isolation_source qualifier in the GenBank entries, were categorised into 18 categories, which were then summarized and presented as <u>Fig. S3</u>.

## Attraction assay

Preliminary results (FileS1) revealed that J2 were attracted to the volatiles of a *Pseudomonas* sp. isolate, which shared its V3V4 region sequence with ASV 108751f28645926db7b461f27b822162. To test the chemical attraction of J2 to the isolate’s volatiles, we carried out attraction assays, following [112]. In 12-well plates (Greiner Bio-One), each well contained a fresh basil root 2 cm fragment and a 10 µl pipette tip containing the tested attractant, to test the attraction towards the liquid in the tip in comparison with the attraction to the root. The tip contained either the whole bacterial filtrate (excluding the bacterium) or one of three size determined fractions (<3kDa, 3-100 kDa, >100 kDa), in nine replicates per treatment. One ml of Pluronic-F127 Tris MES buffer gel (Sigma-Aldrich) containing approximately 50 J2 was added to each well. The number of J2 in each tip was recorded after 24 hours under a dissection microscope (Fig. 7).

The attractants we introduced with the pipette tips were prepared as follows. The isolate was incubated in aquaus beef-extract peptone medium (beef extract 3 g L^−1^ ; peptone 10 g L^−1^; NaCl 5 gL^−1^; [129]) for 48 hours at 37 °C. The culture was filtered using a 0.45 µm syringe filter to obtain the bacterial extracts (whole filtrate). The size fractions were then separated using Amicon Ultra-15 centrifugal filter units with Ultracel-PL membrane, following manufacturer’s instructions. *M. incognita* eggs were sieved from infected tomato roots using a set of mesh #200 sieve on top of mesh #500 sieve (Tyler.S.W), and J2 were hatched with a Baermann tray, as described in [112].

## Supplementary material

**Fig. S1:** Observed-ASV alpha-rarefaction curves for each niche.

**Fig. S2:** Longitudinal descriptions of relative abundances of key taxa and their ASVs.

**Fig. S3**: Isolation sources of 16S rRNA sequences of A/N/*P-Rhizobium, Devosia* and AMB-16S-1323.

**File S1:** Description and results of the preliminary attraction assay of *M. incognita* to the *Pseudomonas* isolate.

## Declarations

### Ethics approval and consent to participate

Not applicable.

### Consent for publication

Not applicable.

### Availability of data and materials

The datasets generated and analysed during the current study are available in the National Center for Biotechnology Information (NCBI) BioProject repository under the accession number PRJNA614519. Data and script are archived as a GitHub release (github: https://github.com/DSASC/yergaliyev2020; Zenodo: DOI: 10.5281/zenodo.3724182).

### Authors’ contributions

TY analysed the data. TY and AS wrote the manuscript. TY, RAS and HD carried out the lab work. HD, RAS and AS collected the samples. AS, SP, DMB and SR, designed the study and edited the manuscript. AS conceived the study. All authors read and approved the final manuscript.

## Acknowledgements

The authors like to thank Dr. David H. Lunt and Dr Keith Davies for very helpful discussions.

