## Supplementary material for "The bacterial community structure dynamics in *Meloidogyne incognita* infected roots and its role in worm-microbiome interactions": File S1

### *Supplementary file 1*: The isolation and identification of *Pseudomonas* sp. *nov.* isolated from RKN infected eggplant roots, and its attractiveness to *M. incognita*.

### Methods

#### Isolation

*Meloidogyne incognita* infested Eggplant roots were washed with ultrapure water, soaked in sodium hypochlorite 1% (v/v) for 5 minutes, and washed with water again. The dissected fragments were placed in 15 ml Phosphate Buffered Saline (PBS; Biological Industries) and stirred vigorously. Forty µl of the PBS were then smeared on a beef-extract-peptone agar (BEPA) plate (Huang et al. 2015), which was then incubated for 72 hours at 37 °C under anoxic conditions, together with a blank BEPA, as control. One colony of each unique form detected was streaked on a fresh BEPA plate and incubated again. Isolates were cultured in 5 ml of liquid beef-extract-peptone medium.

#### 16 rRNA sequencing

To identify the isolates, DNA was extracted from each isolate with the DNeasy blood and tissue DNA extraction kit (Qiagen) and the 16S-rRNA gene was amplified by PCR reaction using the primers BacSSU_FAM27f (5'-GAGTTTGATCMTGGCTCAG-3') and BacSSU_1407R (5'-GACGGGCGGTGTGTRC-3'). The PCR reaction (one cycle at 95 °C – 3 min; 33 cycles at 98 °C – 20 s, 57 °C – 15 s, 72 °C – 21 s; one cycle at 72 °C – 1 min) was carried out in triplicate with the KAPA HiFi HotStart ReadyMix PCR Kit (Kapa Biosystems) following the provided instructions, on a SimpliAmp thermal cycler (ABI). PCR products were purified with Agencourt AMPure XP and directly sequenced on an ABI PRISM BigDye Terminator sequencer. The 1,380 bp long amplification product was subjected to the online blast analysis using the whole blast nucleotide database.

### Attraction assays

Attraction assays were carried out in two 12-well plates with each well comprising 1 ml Pluronic P-127 Tris MES buffer gel, prepared as previously described (Williamson and Čepulytė 2017), and mixed with approximately 200 J2 larvae. The isolate contained by a pipette tip was placed in the well with an eggplant root fragment, 20 mm apart from one another and the number of attracted J2 larvae to each was compared (Fig. 1). This was performed in eight replicates for each bacterial isolate tested. For control, the root fragment was placed in a well together with a pipette tip containing sterile BEPA medium, and attraction was compared as mentioned above. The assay lasted for 10 minutes, after which all J2 larvae reaching the rhizoplane were counted as well as J2 larvae which aggregated in the bacteria-containing pipette tip.

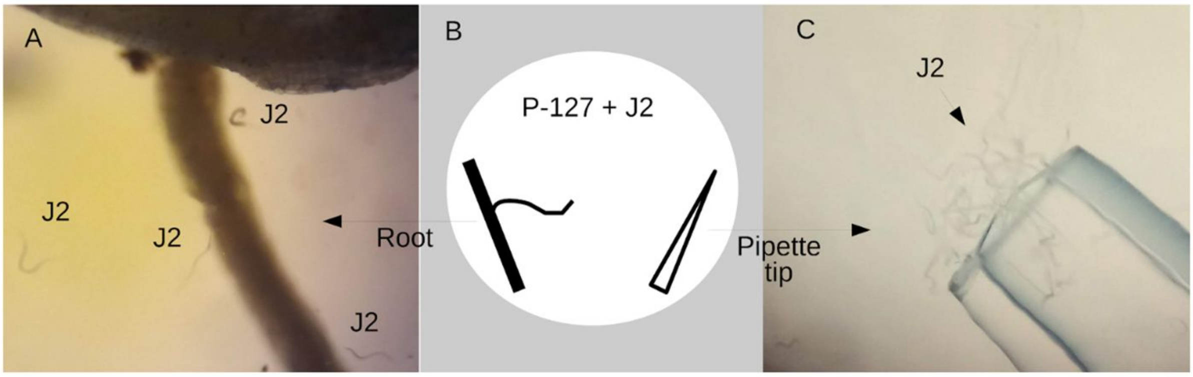


**Fig 1:** Attraction assay setup: Each well (B) contained a root fragment (A) and a pipette tip containing the tested bacterium, bacterial filtrate or sterile medium control. J2: second stage juveniles. P-127: Pluronic P-127 Tris MES buffer gel.

A second attraction assay was designed to compare the attraction of the isolate, with that of its filtrate. To produce a filtrate, the bacterium was cultured in BEPA medium for three days, as described above. The medium was then filtered through a 0.45 micron pore-size filter. The assay was carried out as above, using basil roots, with 4 treatments in 4 replicates each: (i) root vs. the isolate, (ii) root vs. the filtrate, (iii) root vs. the sterile medium and (iv) root only. The assay plate was incubated overnight, after which, the roots were stained following (Thies et al. 2002), and the J2s on the roots and tips were counted using a dissection microscope.

### Results

#### Attraction assays

Six unique colony forms were identified in the isolation process, two of which, isolates 1 and 6, appeared to be bacterial. In the attraction assays, the root fragments attracted 5 to 17 J2 larvae, when co-incubated with isolate 6 or control (containing no bacterial isolates). In all replicates containing isolate 1, more than 70 J2 larvae were attracted to the bacteria-containing tip within the inspected time frame. No J2 larvae were observed to be attracted to a pipette tip containing either isolate 6 or a sterile medium (Table 1).

**Table 1. First attraction assay**

| Trial | Attraction to root mean (min, max) | Attraction to tip mean (min, max) |
|---|---|---|
| Root + isolate 1 | 8.25 (6,11) | 90 (70,120) |
| Root + isolate 6 | 7.25 (5,10) | 0 |
| Root + empty pipette tip | 9.9 (7,13) | 0 |

In the second attraction assay, both the filtrate and the bacterium were at least twice as attractive for the J2s than the root fragment (Table 2).

**Table 2: Second attraction assay**

| Trial | Attraction to root mean (min, max) | Attraction to tip mean (min, max) |
|---|---|---|
| Root + isolate 1 | 8 (4, 11) | 20 (12, 35) |
| Root + isolate 1 filtrate (no bacteria) | 7.33 (5, 7) | 19.25 (1, 66) |
| Root + sterile medium | 9.67 (4, 15) | 9.25 (1, 14) |
| Root only | 22.5 (19, 26) | – |

### Identification of Isolate 1

To determine whether this isolate, which sequence is provided in Table 3, is a novel species, the 318 most similar sequences were retrieved from GenBank using the online version of BLAST. These sequences represented 220 species. The sequences were aligned using MAFFT (Katoh et al. 2002) with 1,000 maximum iterations, and the resulting alignment was trimmed with the gappyout algorithm in trimAl (Capella-Gutiérrez et al. 2009). A phylogenetic tree was reconstructed with FastTree 2.1 (Price et al. 2010), with the GTR substitution model. This data was used to compute a distribution of intra-specific distances (Fig. 2), based on either the proportion of divergent positions (Fig. 2A) or tree-branch distances (Fig. 2B). Both methods revealed the distance between the isolate and its closest sequenced relative (GenBank accession NR_151929.1) is significantly larger than that expected within a species (*P* value <0.0227 and *P* value <0.0045, respectively). Therefore, the newly isolated bacteria is a novel species comprising a SSU sequence which is 1.7% divergent from its closest relative.

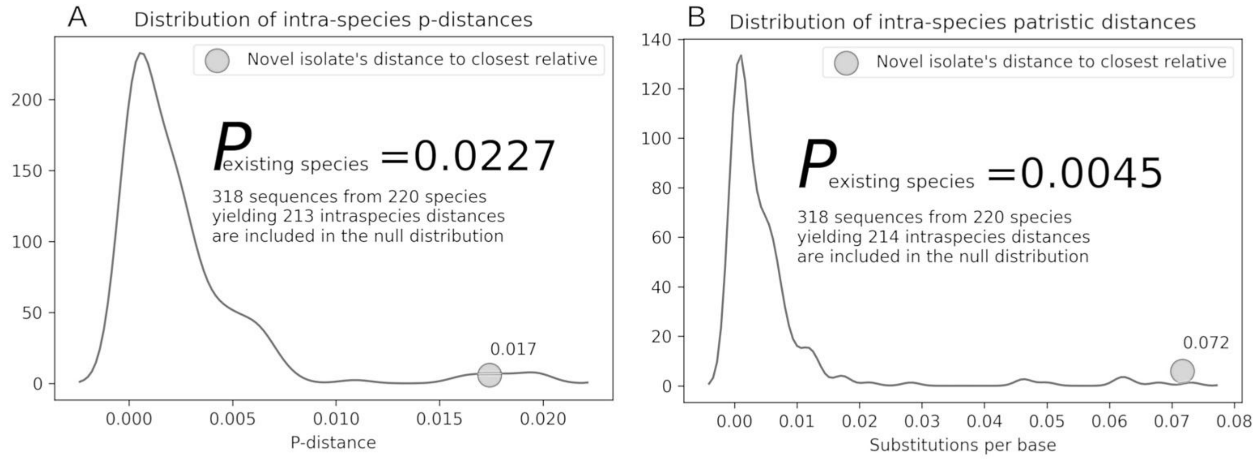


**Fig. 2:** The distribution of intra-specific (A) p-distances and (B) patristic distances. The distance of isolate 1 from its closest sequenced relative is indicated by a gray sphere.

**Table 3. The 16S rRNA sequence of isolate 1**

```
GATAGAGAGGCTGCTGTAGAATGCGCGCCTCGGTTGAGACGAAAGGCTTAACCAACTGTTCTTTAACAACTGAATCAAGCAATTCGTGTGG
GTGCTTGTGAGGTAAGACTGATAGTCAACTGATTATCAGCATCACAAAGCAACACTCGTTAATTCGAGAGTTACCTTTCATTAATTTGAAA
GTTTTGCGATTGCTGAGCCAAGTTTAGGGTTTTCTCAAAACCCAAGCAGTATTGAACTGAAGAGTTTGATCATGGCTCAGATTGAACGCTG
GCGGCAGGCCTAACACATGCAAGTCGAGCGGATGAGAGGAGCTTGCTCCTTGATTTAGCGGCGGACGGGTGAGTAATGCCTAGGAATCTGC
CTGGTAGTGGGGGATAACGTCCGGAAACGGGCGCTAATACCGCATACGTCCTACGGGAGAAAGCAGGGGACCTTCGGGCCTTGCGCTATCA
GATGAGCCTAGGTCGGATTAGCTAGTTGGTGAGGTAATGGCTCACCAAGGCGACGATCCGTAACTGGTCTGAGAGGATGATCAGTCACACT
GGAACTGAGACACGGTCCAGACTCCTACGGGAGGCAGCAGTGGGGAATATTGGACAATGGGCGAAAGCCTGATCCAGCCATGCCGCGTGTG
TGAAGAAGGTCTTCGGATTGTAAAGCACTTTAAGTTGGGAGGAAGGGCATTAACCTAATACGTTAGTGTTTTGACGTTACCGACAGAATAA
GCACCGGCTAACTTCGTGCCAGCAGCCGCGGTAATACGAAGGGTGCAAGCGTTAATCGGAATTACTGGGCGTAAAGCGCGCGTAGGTGGTT
CGTTAAGTTGGATGTGAAAGCCCCGGGCTCAACCTGGGAACTGCATCCAAAACTGGCGAGCTAGAGTACGGTAGAGGGTGGTGGAATTTCC
TGTGTAGCGGTGAAATGCGTAGATATAGGAAGGAACACCAGTGGCGAAGGCGACCACCTGGACTGATACTGACACTGAGGTGCGAAAGCGT
GGGGAGCAAACAGGATTAGATACCCTGGTAGTCCACGCCGTAAACGATGTCAACTAGCCGTTGGGTTCCTTGAGAACTTAGTGGCGCAGCT
AACGCATTAAGTTGACCGCCTGGGGAGTACGGCCGCAAGGTTAAAACTCAAATGAATTGACGGGGGCCCGCACAAGCGGTGGAGCATGTGG
TTTAATTCGAAGCAACGCGAAGAACCTTACCTGGCCTTGACATGCTGAGAACTTTCCAGAGATGGATTGGTGCCTTCGGGAACTCAGACAC
AGGTGCTGCATGGCTGTCGTCAGCTCGTGTCGTGAGATGTTGGGTTAAGTCCC
```
