## Supplementary figures and images for "The bacterial community structure dynamics in *Meloidogyne incognita* infected roots and its role in worm-microbiome interactions"

### Fig. S1

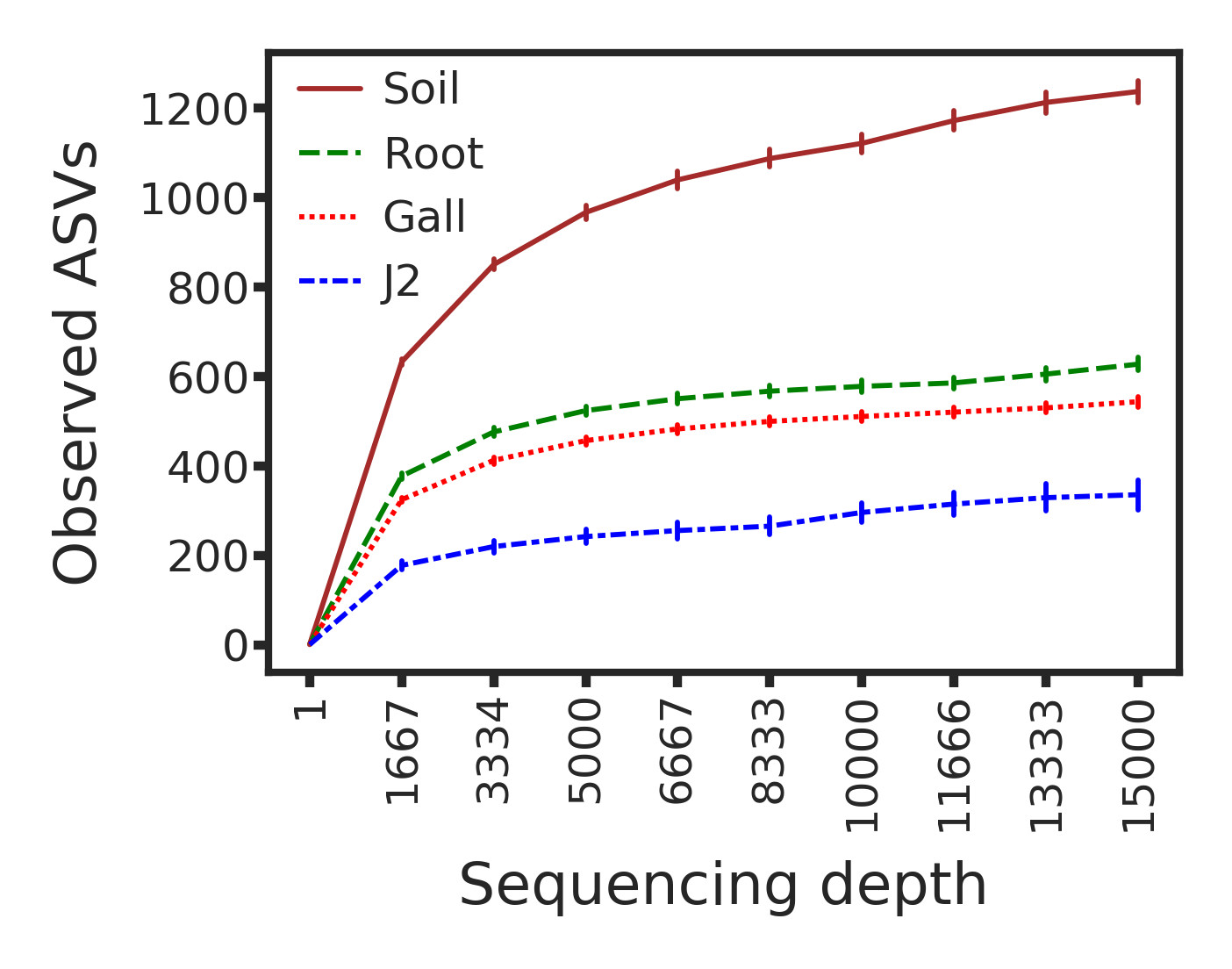

### Fig. S2

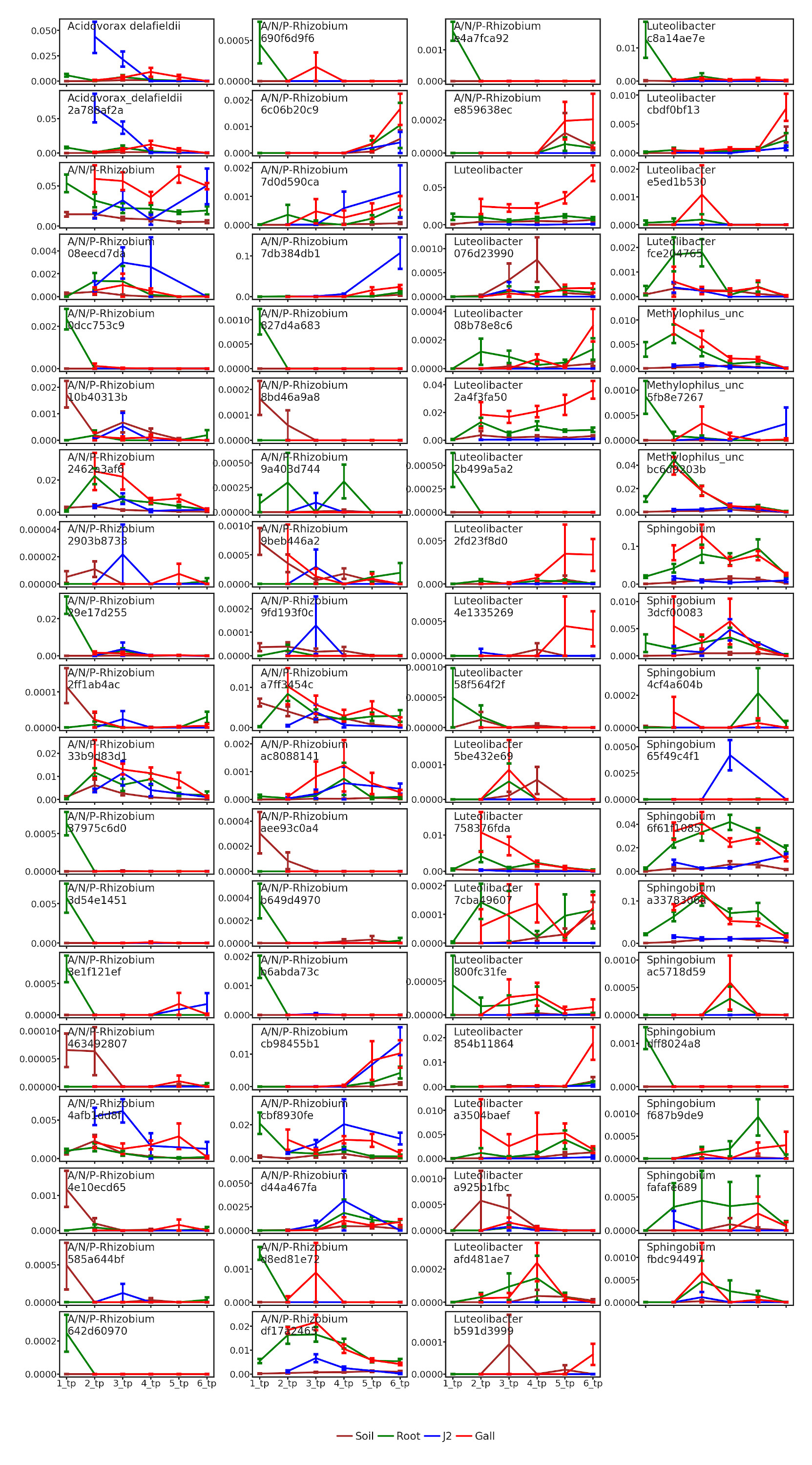

### Fig. S3

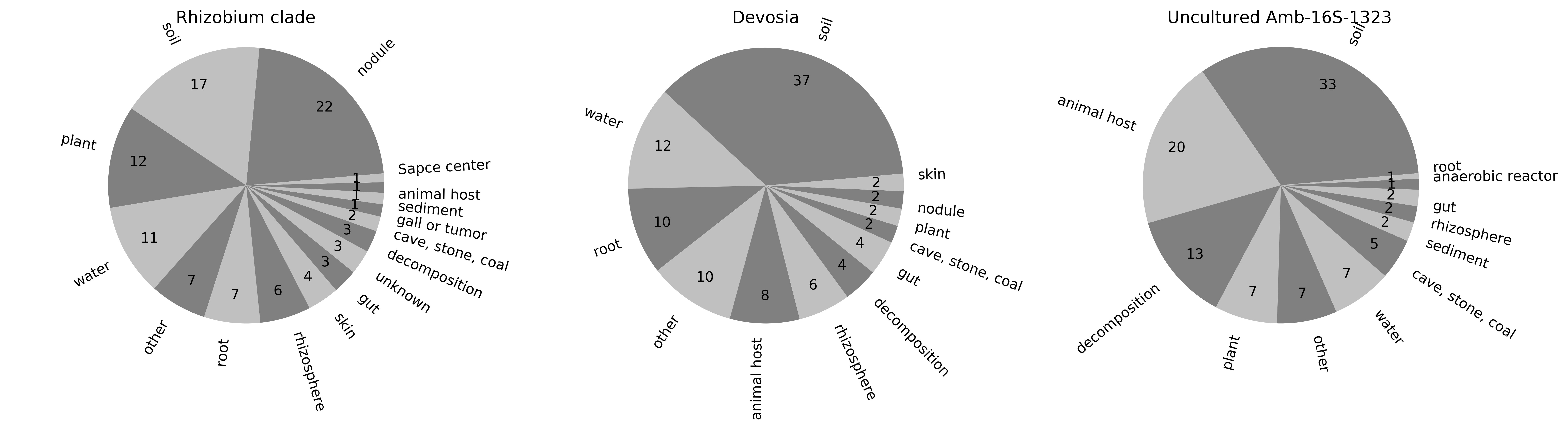
